# Feeding location of aphid prey affects life history traits of a native predator

**DOI:** 10.1101/429415

**Authors:** Ximena Cibils-Stewart, James Nechols, Kristopher Giles, Brian P. McCornack

## Abstract

The green peach aphid (GPA), *Myzus persicae S*., and the cabbage aphid (CA), *Brevicoryne brassicae L*., exhibit a feeding preference when exposed to different canola (*Brassica napus L*.) plant structures. Preference may be associated with the concentration and/or composition of allelochemicals; specifically, glucosinolates (GLS). Furthermore, each aphid species employs a different strategy for tolerating plant defensive chemistry; GPA excretes glucosinolates while CA sequesters these toxic compounds. Given these different detoxification mechanisms, it is possible that both feeding location and aphid species may affect prey suitability for *Hippodamia convergens* larvae. We conducted a factorial laboratory experiment to evaluate whether one or both factors impacted predator fitness. We hypothesized that plant structures with higher GLS concentrations will negatively affect the development and survival of immature predators, which will also vary based on prey detoxification strategies. Results confirm that when predators fed on either aphid species reared on canola structures having lower GLS concentrations, development was faster than when they fed on aphids reared on structures having higher GLS. Although predators consumed more GPA than CA, consumption rates did not differ between rearing location for GPA, but fewer CA were consumed when reared on reproductive canola parts. These findings suggest that: 1) plant-mediated differences in prey quality exist for canola aphids; 2) the type of adaptation used by aphids to overcome plant defenses have important consequences for prey suitability; and 3) reduced feeding by *H. convergens* larvae on unsuitable prey may offset deleterious effects of plant defenses against aphid pests. By evaluating source-sink dynamics at the plant level, we can further understand trophic interactions involving plant defenses and how these interactions may influence community dynamics and structure.

## Introduction

During the past decade, commercial production of winter canola (*Brassica napus L*.) in the Southern Great Plains of the USA has increased substantially (Peeper et al. 2009, Franke et al. 2009, U.S Canola Association; USDA-NASS 2018). Rapid adoption is driven primarily by agronomic and economic benefits provided to producers rotating canola with winter wheat (*Triticuma estivum L*.), which has been a staple crop in this region for decades (Peterson and Westfall 1994, Blackshaw et al. 2001, Giles et al. 2008, Giles and Walker 2009, Peeper et al. 2009, Franke et al. 2009). It is reasonable to assume that spatial and temporal changes in the landscape due to the expansion of novel crops such as winter canola may cause changes in arthropod communities, including different utilization of aphid prey by predators. Depending on the quality of prey available to predators, and on habitat choice offered by these changes in landscape, the outcome on existing predator communities selecting these new prey could be negative (Lu et al. 2012, Wolfenbarger et al. 2008, Landis et al. 2008).

In a canola-wheat rotation, unique complexes of aphid species that are host specific attack each crop. As a result, aphid species found colonizing canola may, or may not, serve as suitable alternate prey for natural enemies that once inhabited a predominantly wheat or grassland landscape (Franke et al. 2009, Knodel 2011, Jessie et al. 2015). Consequently, introducing new prey items into a relatively stable cropping system like wheat, may alter existing ecosystem services by either providing new sources of predators and parasitoids or pulling services away from wheat systems, thus reducing beneficial insects within these changing landscapes. Furthermore, in the context of management, canola landscapes may be classified as “sinks”, since 90% of the canola produced in the Southern Great Plains is treated with broad-spectrum insecticides (i.e., synthetic pyrethroids) annually (Franke et al. 2009). Sink-type habitats are often referred to as “ecological traps”, meaning that they cannot sustain or increase survival and reproduction of beneficial insects and pests (Dwernychuk and Boag 1972). Additionally, aphids attacking canola interact with secondary compounds, glucosinolates (allelochemicals) produced by the canola plant chemical defenses; these interactions may have direct fitness consequences for biological control organisms that prefer these aphids as a resource, thus acting as biological sinks (Cole 1997ab, Francis et al. 2000, 2001, Pratt et al. 2008, Kos et al. 2011a, 2011b, Jessie et al. 2015). In contrast, less than 16% of the winter wheat fields in the same areas are treated with insecticides annually (Giles et al. 2008, Giles and Walker 2009). Despite the continued increase in canola acreage and corresponding increases in insecticide usage, we lack understanding of potential landscape-level impacts of adding canola to a historically wheat-dominated landscape (Brewer et al. 2004, Franke et al. 2009, Knodel 2011). However, to understand the broader landscape-level effects, it is also imperative to focus on plant-level interactions that govern predator-prey dynamics at a local scale (within-plant).

Foliar insecticide applications for aphid management in canola coincide with the flowering stage, which is peak attraction time for pollinators and natural enemies dispersing from nearby habitats and using canola as a resource (Baggen et al. 1999). Consequently, aphid species attacking canola during flowering include a generalist, the green peach aphid or GPA (*Myzus persicae*) and a specialist, the cabbage aphid or CA (*Brevicoryne brassicae*) (Franke et al. 2009, Berlandier et al. 2010, Royer and Giles 2008, Royer and Giles 2017). Green peach aphid is a polyphagous pest that can utilize over 400 plant families (van Emden at al. 1969), being able to mitigate glucosinolates produced by canola through excretion (Weber 1985, Pratt et al. 2008, Hopkins et al. 2009). In contrast, <u>CA</u> is a perennial pest that is restricted to plants in the Brassicaceae family, which includes winter canola (Hughes 1963). This specialist species has evolved mechanisms to sequester glucosinolates using an aphid-endogenous enzyme that detoxifies these secondary plant compounds, thus acquiring the compound and becoming itself toxic to natural enemies (Weber 1985, Kazana et al. 2007, Pratt et al. 2008, Hopkins et al. 2009, Jessie et al. 2015). Under field conditions, these two species have different vertical distributions within the canola canopy (Cibils et al. 2015). Specifically, GPA are observed predominantly on the lower leaves (vegetative tissue) of the plant, while CA are observed primarily colonizing the flowering racemes (reproductive tissue). Reasons for these differences may be the result of top-down (predator preference) or bottom-up forces (plant quality or nutrient composition), competition between aphid species (space, resources, etc.), or a combination of factors (Merritt 1996, Idris and Roff 2002, Smallegange et al. 2007, Pekar 2005, Berberet et al. 2009, Fernandes et al. 2011, Cibils-Stewart et al. 2015). Bottom-up forces, such as variation in host quality (i.e., allelochemicals) influence prey quality and toxicity (fatty acid content or toxicity through sequestration), which has direct repercussions on mortality, development, growth rates, fecundity and recruitment of natural enemies in other brassica species (Gabrys et al 1997, Cole. 1997a, Giles at al. 2002, Brown et al. 2003b, Giles et al. 2005, Lambton and Hassall 2005, Smallengange et al. 2007, Van Dam et al 2008a, 2008b, Guigo et al. 2010, Kabouw et al 2010, Kramer et al. 2011). Consequently, for natural enemies using canola aphids as a resource, prey/host selection can directly affect fitness (Idris and Roff 2002, Pekar 2005, Berberet et al. 2009, Fernandes et al. 2011, Jessie et al. 2015). For instance, Jessie et al. (2015) demonstrated reduced fitness of insect predators when feeding on brassicae specialist canola aphids (CA) that sequestered higher glucosinolates concentrations; however, aphids were collected from uniform infestations of seedling canola plants and may not reflect prey quality differences as influenced by typical within-plant distributions (Cibils-Stewart et al. 2015). To our knowledge, these plant-level interactions between aphids occupying different structures, and natural enemies that encounter such prey patches have not been studied for canola specifically.

A complex of natural enemies is observed annually on winter wheat in the Southern Great Plains and the impacts on cereal aphid suppression are well documented (Chambers 1986, Michels et al. 2001). Lady beetles (Coccinellidae), green lacewings (*Chrysopera* spp), damsel bugs (*Nabis* spp), syrphid flies (*Syrphidae* spp), and spiders (Araneae), among others are abundant in these wheat-dominated landscapes (Chambers 1986, Michels et al. 2001); these natural enemies are also abundant in winter canola (Cibils-Stewart 2013). Lady beetles specifically play a critical role in cereal aphid suppression (Kring et al. 1985, Michels and Behle, 1991, Obrycki and Kring 1998, Michels at al. 2001, Phoofolo et al. 2007, Obrycki et al. 2009) and also co-occur with canola flowering (Jessie 2017). Among all coccinellid species active in spring, which is when canola and wheat aphids are most abundant, the convergent lady beetle, *Hippodamia convergens*, is the most dominant coccinellid species (Nielson et al. 1959, Dogan et al. 1996, Nechols and Harvey 1998, Michels et al. 2001, Michaud and Qureshi 2006, Jessie 2017). Similar to other predatory lady beetles, *H. convergens* are effective predators both as adults and larvae and are capable of using alternative foods such as pollen, sap, or nectar when prey numbers are low during the adult life stage (Hodek 1973, Lundgren 2011); these alternative resources become readily abundant during early spring with the addition of flowering canola in the landscape. While adult lady beetles have a high dispersal capacity that enables active foraging of available resources in wheat and surrounding landscapes (Hodek 1973, Hajek 2004, Seagraves 2009), immature stages are more restricted to maternally selected prey patches at a local scale or within-field level (Evans 2003). Therefore, gravid females can indirectly affect larval fitness through searching and oviposition decisions (Evans 2003). Consequently, recruitment plays an integral role in determining the impact of adding new crops with novel prey complexes to existing predator-prey systems. The addition of novel prey found in canola, plus the surplus of pollen and nectar resources provided by canola, may directly influence adult lady beetles oviposition decisions. Hence, in the context of source-sink relationships, recruitment can play an integral part in determining the impact of adding new resources to predator-prey system. Thus, life history traits of immature lady beetles feeding on these novel prey patches may be affected directly as a consequence of maternal oviposition. Since this complex of aphids also interacts with canola secondary compounds in different ways, the effect of adding novel prey to existing predator services within the landscape is not known.

To our knowledge, the suitability of canola aphids reared on different canola plant structures as a suitable diet for immature *H. convergens* has not yet been evaluated. Although different prey species can be accepted by predatory lady beetles, suitability levels across aphid species in canola-wheat landscapes needs further study. More specifically, suitability determines if aphid prey are essential or alternative prey for lady beetles (Hodek and Honek 1996, 2001). While essential prey ensures survival and fecundity of predators, alternative prey only provides nutrients necessary for survival (Hodek and Honek 1996, Kalushkov and Hodek 2001). The objectives of this study were to compare various canola aphid diets on immature *H. convergens* life history traits, which included survival, development, prey consumption rates, pupal weight, and adult ecolsion times. Aphid diets were accomplished by rearing glucosinolate-excreting (GPA) or glucosinolate-sequestering (CA) aphids on either reproductive or vegetative canola plant tissues. Two additional control diets were used: the soybean aphid, *Aphis glycines* (Matsumura), that serve as a non-canola aphid prey, and an artificial diet, the non-aphid diet mix comprised of eggs of the Mediterranean meal moth, *Ephestia kuehniella* (Zeller).

We predicted that: 1) aphid prey reared on reproductive parts of canola plants would be less suitable for *H. convergens*; and 2) independent of aphid feeding location, aphids sequestering glucosinolates (i.e., CA) would be the least suitable diet (in terms of fitness measurements) compared to aphids that excreted glucosinolates (i.e., CA). By evaluating source-sink dynamics at the plant level, we can further understand trophic interactions occurring between species (both pest and beneficial) and potentially predict landscape-level processes. Potentially, such interactions may influence insect community dynamics and structure within wheat-canola agroecosystems (Guigo and Corff 2010).

## Materials and Methods

### Aphids and plants

Canola aphids (GPA and CA) used in feeding assays were collected from natural field populations in Barber County Kansas (N 36.998414, W –98.456797) in April 2011, and maintained under laboratory conditions at Kansas State University (Manhattan, KS). Aphids were transported to the laboratory in coolers, where they were provided with young, potted canola seedlings as a food source. Aphid colonies were maintained in growth chambers (Bio-Temp Scientific Inc., BT-1–49 WC, Bradenton, FL) at 22 ± 2 ºC, 60–70% RH, with a 16:8 hr (light: dark) photoperiod (Kos et al. 2011a). Canola (cv. Riley) was seeded in a special soil mix that contained sulfur, ammonium nitrate, micronutrients, peat moss, perlite, and two types of slow release fertilizers (proprietary soil blend, M. Stamm, canola breeder, Kansas State University and Oklahoma State University). Plants were maintained in the greenhouse at 22 ± 3ºC under natural light during daylight hours and supplemented with artificial lights at night; photoperiod was kept at 16:8 hr (light: dark). Canola plants were artificially vernalized in a growth chamber for approx. 2 months at 12:12 hr (light: dark) and constant 4ºC to induce reproductive maturity (Murphy and Scarth 1994). Vernalized plants were then used for no-choice experiments.

Colonies of *Aphis glycines* (SA, biotype 1) were obtained from a source colony maintained at Iowa State University (Ames, Iowa). SA were reared at 22 ± 2ºC, 60–70% RH, 16:18hr (light: dark) photoperiod in a growth chamber (Bio-Temp Scientific Inc., BT-1–49 WC, Bradenton, FL) on susceptible soybean plants (cv. A1026832, Asgrow®, St Louis, MO) grown in horticultural flats (5 × 10 cells) (Deepots, Hummert International). Flats were placed in plastic storage tubs (587.7 × 42.9 × 16.2 cm, L×W×H) (Sterilite storage box, Leominister, MA) filled with water; tubs allowed pots to receive water *ad libitum*. Once soybean plants were in vegetative stages 3 to 6 (V3-V6) (Fehr et al. 1971), soybean aphids were transferred to plants. Two flats of soybeans were planted in the greenhouse bi-weekly to adequately maintain soybean aphid colonies. SA was included in these experiment as non-canola prey and served as a control (i.e., non-GLS aphid diet).

### Lady beetles

*Hippodamia convergens* adults were collected from commercial cornfields in Washington County Kansas in August 2012 (N 39.999561, W –97.344964). Adult beetles were collected (*n* = 300) using a hand-held aspirator and brought to Kansas State University (Manhattan, KS) in coolers. Adults were maintained at 22 ± 2ºC, 60–70% RH, 16:8 hr (light: dark) photoperiod in a growth chamber (Percival, AR22L, Perry, IA). *H. convergens* adults were provided with honey and a supplementary, artificial diet mix that consisted of a mixture pollen substitute, tropical fish flakes, cichlid pellets, sundried *Gammarus pulex* and *Ephestia kuehniella* eggs *ad libitum;* non-aphid diet mix (see Lundgren et al. 2011 for complete diet composition). *E. kuehniella* eggs were obtained from a commercial supplier (Beneficial Insectary Inc., Redding, CA). Additionally, SA from laboratory colonies (described above) were included as part of the daily *H*. *convergens* diet to avoid reproductive diapause (Vargas et al. 2012). *H. convergens* adults were kept in commercial rearing cages (61 × 61 × 61cm) (BugDorms, BioQuip Inc., CA, USA) and cleaned weekly.

All adults were left in cages for 2 wk to facilitate multiple mating events and increase chances of viable egg production. Adults were sexed using size, morphological characters on the last abdominal segment, and overall coloration (Kova’r 1996). Once sexed, females were placed in individual plastic containers (4.5 × 4.5 × 1.5 cm) and provided with SA *ad libitum*, a water-saturated cotton gauze, a droplet of honey (0.3 × 0.2 cm diameter), and approx. 0.5 mg of non-aphid diet mix daily. Individual containers were checked daily for egg production. Once eggs were deposited, females were moved to new containers and eggs remained undisturbed in the same container until hatching. Newly hatched larvae were placed into individual cells (128 per tray) of a clear bioassay tray (BAC128, Bio-Serv®, Frenchtown, NJ). Larvae were restricted to the same cells using clear, 16-cell adhesive lids (BACV16, Bio-Serv®, Frenchtown, NJ). Larvae were provided with *E. kuehniella* eggs *ad libitum* and water-saturated cotton gauze throughout their larval development. After rearing one generation of *H*. *convergens* in the laboratory from field-collected adults, newly hatched larvae from this first generation (i.e., second generation larvae) were used for all no-choice bioassay trials. To avoid maternal effects, larvae of multiple mothers were randomly selected. Colonies were maintained for future studies; field collected female and male adults were added regularly to avoid consanguinity or inbreeding effects (Soares et al. 2001). A total of one hundred sixty-eight immature lady beetles (28 individuals per diet treatment), from eggs of 50 adult females, were evaluated to determine if prey feeding location on canola influenced immature life history traits of predators.

### No-choice bioassay

We tested effects of various prey diets on the life history attributes of immature *H*. *convergens* using a no-choice feeding assay. Different aphid diets were produced by restricting *B. brassicae* or *M. persicae* on either reproductive (flowering racemes) or vegetative (mid-canopy leaves) canola plant structures using enclosure cages (Soper et al. 2012, Cibils-Stewart et al. 2015). In companion experiments, Cibils-Stewart (2013) showed a lack of cage effect under field and greenhouse conditions across multiple years using the same cage design. Cages (23 cm diameter, 71 cm length) were placed on the same plant to minimize individual plant affects (e.g., plant size, nutritional composition, etc.) on aphid responses. Each cage consisted of white, no-see-um mesh (Quest Outfitters, Sarasota, FL) with zippered tops; zippers provided access to either the flowering raceme or the vegetative leaf after a cage was secured to a plant structure. The base of each enclosure cage was secured to the canola plant using 15 cm plastic cable-ties (Gardner Bender, Butler, WI), which were located below the last flower of the flowering raceme, or at the node between the leaf and the plant stem. To allow free-movement of aphids within the cage, cylindrical supports made of 14-gauge, galvanized steel wire (Impex Systems Group, Inc. Miami, FL) were added to each cage. These wire supports kept the mesh from resting on the flowers or the leaves or disrupting growing aphid populations. Canola plants used to produce aphid treatments were arranged in a completely randomized design in the greenhouse at 22 ± 3ºC, under natural light supplemented by artificial lights in a 16:8 hr (light: dark) photoperiod. Adult aphids (5 per cage) were placed on plant structures 3 wk prior to initiating the experiment to ensure enough aphid for each diet treatment.

Newly hatched (< 8 hours old), second-generation *H*. *convergens* larvae were selected and individually placed in bioassay tray cells; larvae were still feeding on their egg cases prior to initiating the bioassay. Larvae were assigned to one of six diets: *M. persicae* restricted to feeding on reproductive (GPA-R) or vegetative (GPA-V) canola tissues, *B. brassicae* restricted to feeding on reproductive (CA-R) or vegetative (CA-V) canola tissues, non-canola aphid-control (*Aphis glycines* [SA]), and non-aphid diet (*E. kuehniella* eggs). Larvae were fed the same diet for the duration of their larval development. Aphids were collected daily from infested plants and used for the feeding assays to provide larvae *ad libitum* prey; *ad libitum* levels incorporate both preference and satiation typical of natural interactions in the field (unpublished data, X.C.S., Giles et al. 2000). Aphids were placed within cells with a fragment of leaf material from their host plants to reduce mortality caused by handling; cells were arranged in a completely randomized design. For the *E. kuehniella* diet, eggs (approx. 0.12 g) were added to cells as needed daily to ensure *ad libitum* supply. All larvae were reared in a growth chamber (Percival, AR22L, Perry, IA) at 22 ± 2ºC, 60–70% RH, 16:8 hr (light: dark) photoperiod.

The main effect of aphid feeding location (reproductive versus vegetative structures) on *H*. *convergens* mortality, development (hatch to adult emergence), and consumption rates were determined through daily observations. For all aphid diets, consumptive effects were further categorized into total number aphids consumed or partially consumed, as well as number of unconsumed aphids. Consumption values were not recorded for *H. convergens* reared on *E. kuehniella* eggs as these larvae were fed eggs *ad libitum*. Larval development (instar change) was determined by presence of exuviae in the bioassay cells. Predator exuviae, any remaining aphids, and plant material were removed daily from all cells and replaced with new diet. Bioassay trays were replaced with new ones weekly to reduce infection and mold growth. After larvae completed development, individual pupal weights were recorded (10^−4^ mg, Denver Instrument, Pinnacle Analytical Scale, Denver, CO) and compared among diet treatments. Pupal mass was used as a measure of adult fitness. Pupae were then placed back into the bioassay trays and days to adult emergence was recorded. Response variables included larvae mortality, development, daily consumption, larval weight, pupal weight, and adult eclosion. The experiment was repeated at three different times (blocks): block 1 had 4 larvae per treatment; and blocks 2 and 3 each had 12 larvae per treatment. Number of replications varied among blocks because of the criteria of selection of larvae used; only newly hatched (< 8 hours old) larvae still feeding on their egg cases were used. Voucher specimens for H. *convergens, B. brassicae, M. persicae*, and *A. glycines* nymphs and adult were deposited in the Kansas State University Museum of Entomological and Prairie Arthropod Research (voucher number 228).

### Statistical analysis

For mortality results, the Area Under the Disease Progression Curve (AUDPC) (Jeger & Viljanen-Rollinson. 2001) method was used to determine differences in mortality among treatments. AUDPC is a method used in plant pathology to quantify disease progress; this method allows for the detection of minimum differences between mortality curves among and between treatments using mean mortality data from the three runs. Although the values provided by the AUDPC are without units, these values were statistically compared using a bidirectional ANOVA (PROC MIXED, SAS 2009); thus, the LS MEANS statement was used to make comparisons between values. The means were separated using the adjusted Tukey method (α = 0.05).

Additionally, all data pertaining to mortality, development, consumption, pupal mass, and of adult emergence of immature lady beetles were analyzed using SAS version 9.3 for Windows (SAS Institute 2009). Assumptions of normality for data were also tested according to the Shapiro-Wilk test statistic (PROC UNIVARIATE, SAS 2009). An analysis of variance (*PROC MIXED*) procedure was performed with diet as a fixed effect and block as a random effect for all parameters. The *LS MEANS* statement with an adjusted Tukey method was used to make treatment comparisons (α = 0.05). Moreover, using only canola aphid diets (CA-R, CA-V, GPA-R, and GPA-V) an analysis of variance (*PROC MIXED*) procedure was performed to test the main effects of previous feeding location of prey on canola, prey species, and the two-way interaction between previous feeding location and aphid species on predator: larval (from egg hatch to pupa) and preimaginal development (from egg hatch to adult emergence), overall consumption, and pupal weights. Mean differences were compared using the least-squares mean difference (LSD) from an adjusted Fisher’s Protected LSD method for multiple comparisons with the significance level of α = 0.05.

## Results

### Mortality

Of the 168 eggs collected from 50 adult females, 101 (60%) survived to pupation, 39 (23%) escaped during early instar stages, and 28 (17%) died. There were no statistical differences among diet treatments in terms of the number of larvae that escaped (*P* =0.38; *F*= 1.15; df = 5, 12), or survived (*P* =0.54; *F*= 0.82; df = 5, 12). Moreover, for those lady beetles that died (17%), the AUDPC values indicated no statistical significances in the time course of mortality among treatments (*P* =0.39; *F*= 1.13; df = 5, 12).

### Larval development

Aphid diet significantly affected larval development (*P* <0.0001; *F*= 25.67; df = 5, 95). Mean larval development (egg hatch to pupation) was shortest for *H*. *convergens* that fed on *Ephestia* eggs (E-EGG), followed by GPA reared on vegetative canola structures (GPA-V), and longest for lady beetles reared on the soybean aphids (SA) and the other canola aphid diets (CA-R, CA-V, and GPA-R) (Fig. 1A). When comparisons of lady beetle larval development were made among the four canola aphid diets excluding controls (CA-R, CA-V, GPA-R, and GPA-V), canola aphid species, aphid feeding location, and the aphid species by feeding location interaction all had significant effect on larval development (Table 1A). Again, larval development was fastest when ladybeetles were on the GPA-V diet treatment, compared to the other canola aphid diet treatments (Fig 2A).

**Table 1.** Results from a mixed model analysis of variance (ANOVA) for *H. convergens* A) larval development (egg hatch-pupae); B) preimaginal development (egg hatch-adult eclosion); daily C) consumed, D) partially consumed, E) unconsumed canola aphids; and F) pupal weights, when exposed to canola aphid diet treatments. Model considers the main effects of 1) prey species (**spp**: *Brevicoryne brassicae* (CA) and *Myzus persicae* (GPA)), and 2) previous feeding location of prey on canola (**loc**: reproductive (R) or vegetative (V) canola plant structures), and their two-way interaction (**spp*loc**). Significant main effects and/or interactions (*P*< 0.05) using Fisher’s Protected LSD, displayed in bold.

| A) | Factor | DF | *F* | *P* |
| --- | --- | --- | --- | --- |
| larval Dev. | spp | 1,71 | 4.56 | **0.0362** |
|  | loc | 1,71 | 8.63 | **0.0045** |
|  | spp*loc | 1,71 | 4.15 | **0.0454** |

| B) | Factor | DF | *F* | *P* |
| --- | --- | --- | --- | --- |
| Preimaginal Dev. | spp | 1,71 | 3.11 | **0.0521** |
|  | loc | 1,71 | 6.45 | **0.0132** |
|  | spp*loc | 1,71 | 70.2 | **0.0008** |

| C) | Factor | DF | *F* | *P* |
| --- | --- | --- | --- | --- |
| Consumed | spp | 1,77 | 97.04 | **<.0001** |
|  | loc | 1,77 | 30.81 | **<.0001** |
|  | spp*loc | 1,77 | 12.79 | **0.0006** |

| D) | Factor | DF | *F* | *P* |
| --- | --- | --- | --- | --- |
| Partially consumed | spp | 1,77 | 165.01 | **<.0001** |
|  | loc | 1,77 | 44.01 | **<.0001** |
|  | spp*loc | 1,77 | 39.44 | **<.0001** |

| E) | Factor | DF | *F* | *P* |
| --- | --- | --- | --- | --- |
| Unconsumed | spp | 1,77 | 9.69 | **0.0026** |
|  | loc | 1,77 | 0.54 | 0.466 |
|  | spp*loc | 1,77 | 2.95 | 0.0901 |

| F) | Factor | DF | *F* | *P* |
| --- | --- | --- | --- | --- |
| Pupal w. | spp | 1,72 | 4.74 | **0.0327** |
|  | loc | 1,72 | 3.63 | 0.0608 |
|  | spp*loc | 1,72 | 0.04 | 0.8508 |

**Figure 1.**
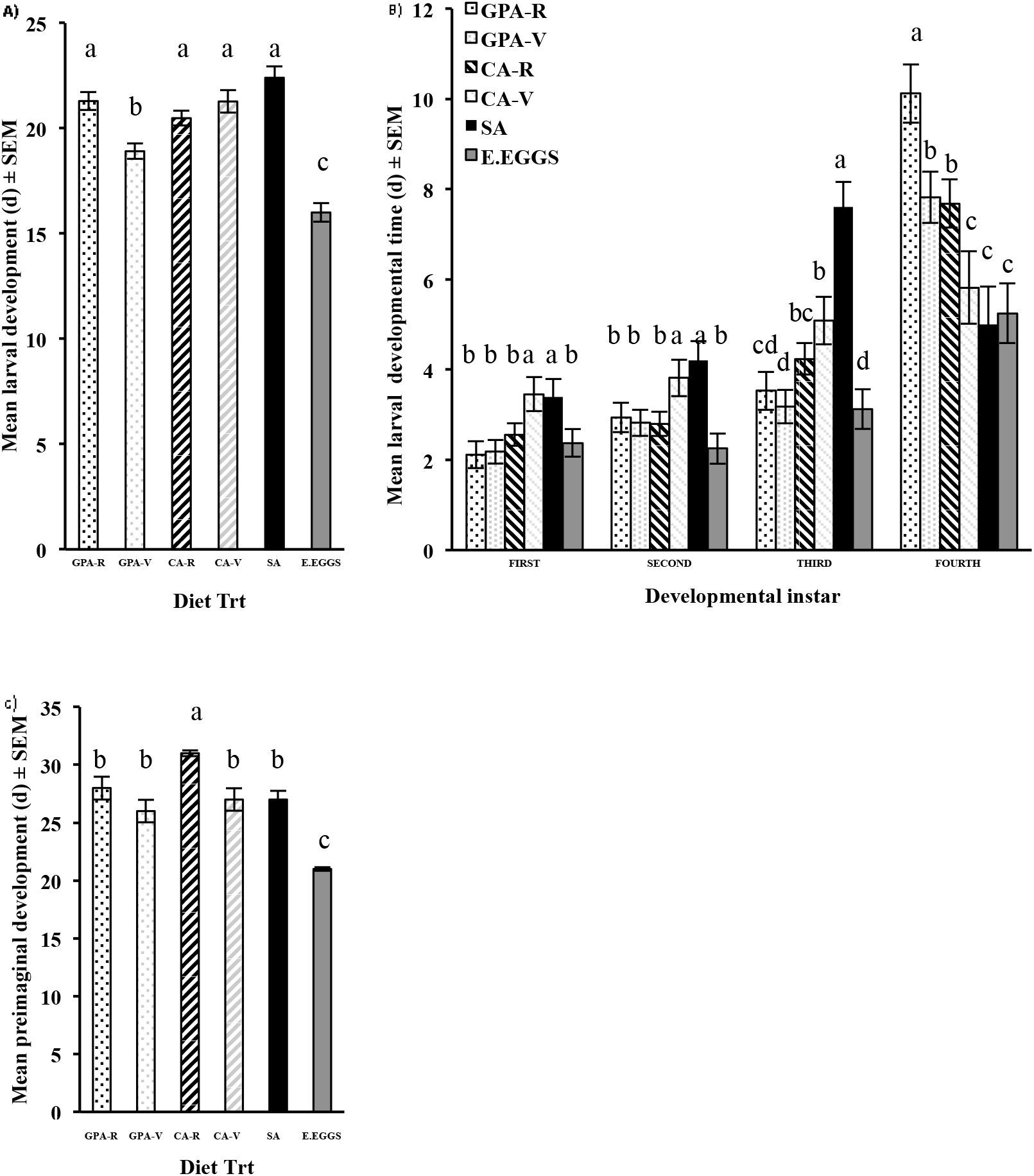
No-choice bioassay assessing effects of different aphid diets on *Hippodamia convergens* A) mean larval (egg hatch to pupation) developmental time (d) ± SEM, B) mean duration of specific instar stages (d) ± SEM, and C) mean preimaginal (egg hatch to adult eclosion) developmental time (d) ± SEM. Diet treatments included: *Myzus persicae* (GPA) and *Brevicoryne brassicae* (CA) previously confined to feeding on reproductive (GPA-R and CA-R) or vegetative (GPA-V and CA-V) structures of canola plant, a non-canola aphid control (*Aphis glycines*, SA), and a non-aphid diet (*Ephestia kuehniella* eggs). Bars with the same letters are not significantly different (*P*< 0.05) using Fisher’s Protected LSD.

When larval development was compared among diets for each larval instar, larvae reared on SA and on CA-V had the longest developmental times during the first two instars compared to the other treatments (first: *P* =0.0134; *F*= 2.98; df= 5, 95, second: *P* =0.0052; *F*= 3.59; df= 5, 95); interestingly, on these treatments larvae spent the least amount of time as fourth instar combined with the E-EGG diet (fourth: *P* <0.0001, *F*= 8.2; df= 5, 95) (Fig. 1B). Furthermore, during the third instar, *H. convergens* larvae restricted to the SA diet had the longest developmental time, followed by the *B. brassicae* diet and the *M. persicae* diet regardless of previous aphid feeding location, and lastly the E-EGG diet on which *H. convergens* larvae developed the fastest (third: *P*<0.0001; *F*= 11.21; df= 5, 95) (Fig. 2B). Consequently, the effect of prey species and prey plant feeding location on *H. convergens* larval development differed among instars. When fed GPA-R, lady beetle larval development was delayed, but only during the last instar (Fig. 1B). However, when *H. convergens* fed on CA-R, longer development occurred only during the four instar when this diet was compared to CA-V (Fig. 1B). Interestingly, prey species and plant feeding location also appeared to affect the relative amount of time lady beetle larvae spent in different instars. For example, the percent of total larval development time (measured in days) spent in the fourth instar (developmental stage where maximum prey consumption is observed) was 24.7 (SA), 32 (CA-R), 44.5 (CA-V), 48.8 (GPA-V), and 54.1% (GPA-R) for each diet, respectively. Interestingly, on the non-aphid diet (E-EGGS) ladybeetles spent 40.3% of their larval development as fourth instars.

### Preimaginal development (egg hatch to adult eclosion)

Time to adult emergence differed among diets (*F*= 15.03; *df*= 5, 95; *P* <0.0001). Emergence took longer for larvae reared on CA-R compared to the other aphid diets, and shorter for ladybeetles on the non-aphid *Ephestia* egg diet (Fig. 1C). Furthermore, when focusing on lady beetle preimaginal development for canola aphid diets only (CA-R, CA-V, GPA-R, GPA-V), significant interactions were observed between previous feeding location of prey in canola, aphid species, and their two-way interaction (Table 1B). *H. convergens* immature exposed to GPA-R diet displayed the longest preimaginal developmental time when compared to the other canola aphid diet treatments (Fig. 2B).

**Figure 2.**
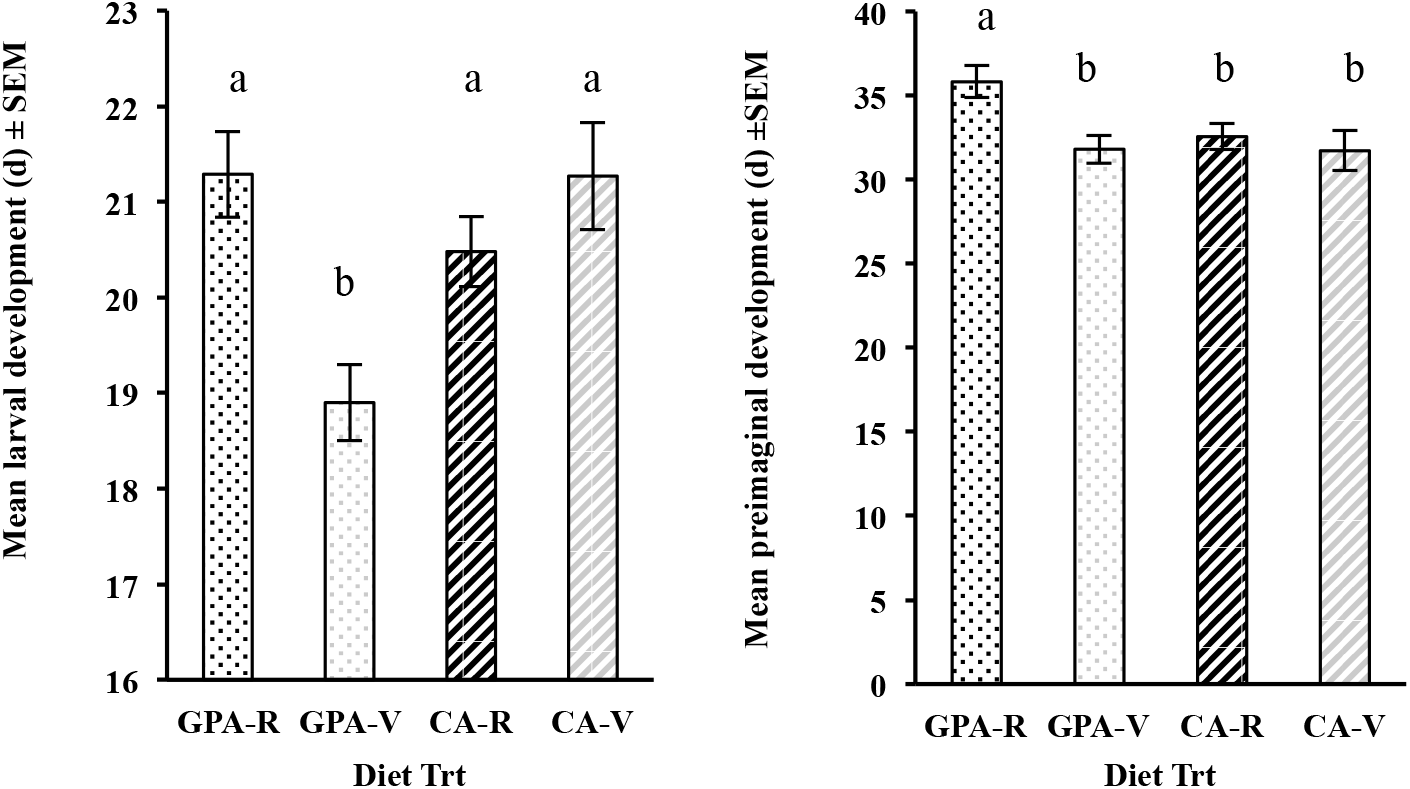
No-choice bioassay assessing effects of different canola aphid diets on *Hippodamia convergens* A) mean larval (egg hatch to pupation) developmental time (d) ± SEM, and B) mean preimaginal (egg hatch to adult eclosion) developmental time (d) ± SEM. Diet treatments included: *Myzus persicae* (GPA) and *Brevicoryne brassicae* (CA) previously confined to feeding on reproductive (GPA-R and CA-R) or vegetative (GPA-V and CA-V) structures of canola plant. Bars with the same letters are not significantly different (*P*< 0.05) using Fisher’s Protected LSD.

### Prey consumption

Aphid daily consumption varied significantly across the diets (consumed: *P* <0.0001; *F*= 48.69; df= 4, 85). The fewest fully consumed aphids were observed on the CA-R diet (Fig. 3A). This treatment combination also resulted in the most partially consumed (Fig. 3B) and unconsumed aphids (Fig. 3C) (partially consumed: *F*= 73.45, df= 4, 85; *P* <0.0001; unconsumed aphids: *F*= 3.85; df= 4, 85, respectively). Interestingly, *H. convergens* larvae consumed more soybean aphids than any other aphid diet treatment (Fig. 3A). Larvae consumed significantly more *M. persicae* than *B. brassicae* regardless of previous aphid feeding location. However, when offered *B. brassicae*, significantly more aphids previously restricted to vegetative structures (CA-V) were consumed than aphids previously restricted to reproductive plant parts (CA-R) (Fig. 3A). Additionally, *H. convergens* larvae left more partially consumed aphids on the CA-R diet, followed by the CA-V diet. In contrast, *M. persicae* previous feeding location (GPA-R, GPA-V) had no effect (*F*= 73.45, df= 4, 85; *P* <0.0001, Fig. 2B). The SA diet had the fewest aphids left partially or unconsumed overall (Fig. 3B-C). Lady beetles on the SA diet consumed 31.4 % more aphids than those exposed to the CA-R (Fig. 3A). On the other hand, lady beetles that fed on the CA-R diet left 93.3% and 69.8% more aphids unconsumed and partially consumed respectively, when compared to those exposed to the SA control diet (Fig. 3B-C).

**Figure 3.**
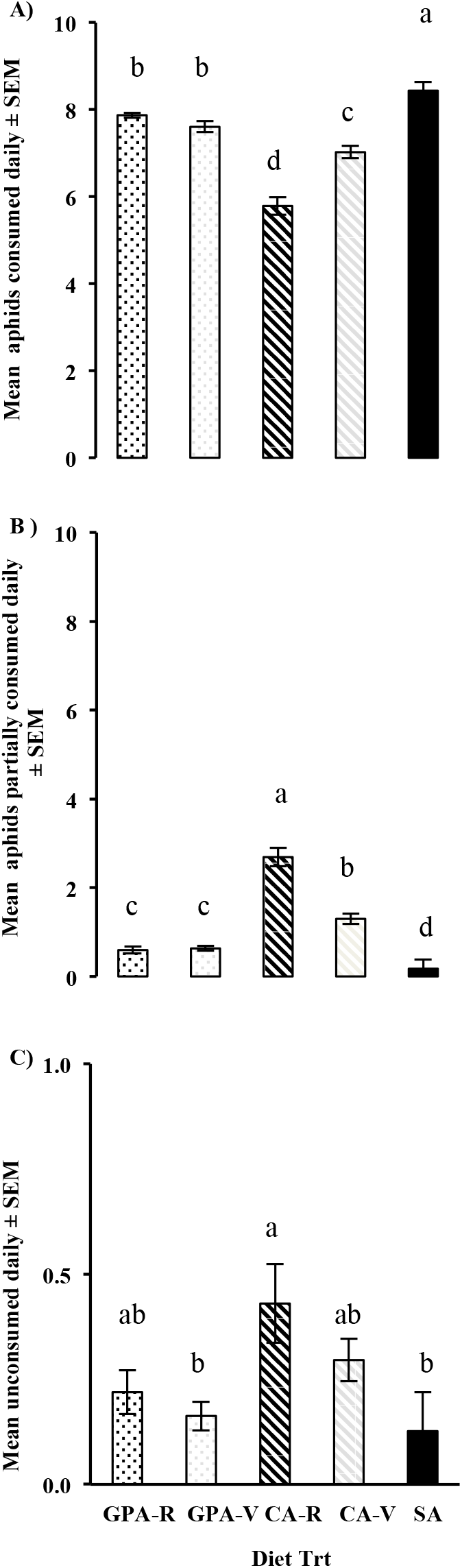
Mean aphid daily consumption ± SEM for *H. convergens* larvae restricted to different aphid diets: *Myzus persicae* (GPA) and *Brevicoryne brassicae* (CA) previously confined to reproductive (GPA-R and CA-R) or vegetative (GPA-V and CAV) structures of canola plants, and an aphid control (*Aphis glycines*, SA). Consumption was categorized as mean number of aphids (10 initial aphids) that were A) fully consumed, B) partially consumed, C) and unconsumed. Bars with the same letters are not significantly different (*P*< 0.05) using Fisher’s Protected LSD.

Comparisons of lady beetle consumption exclusively on canola aphid diets (CA-R, CA-V, GPA-R, GPA-V) showed that the main effects of aphid species (spp) and aphid feeding location (loc), as well as aphid species by feeding location interaction (spp*loc) all had significant effects (Table 1C-E). The greatest consumption was observed on the CA-V and GPA-R diets with the fewest aphids consumed on the CA-R diet (Fig 4A). In terms of partially consumed (Fig 4B) and unconsumed (Fig 4C) canola aphids, CA-R was the least preferred diet, followed by CA-V that only had significant interactions between CA-R in terms of partially consumed aphids. GPA diets, regardless of previous aphid feeding location, were the diets with the least amount of aphids partially or un-consumed (Fig 4C-D).

**Figure 4.**
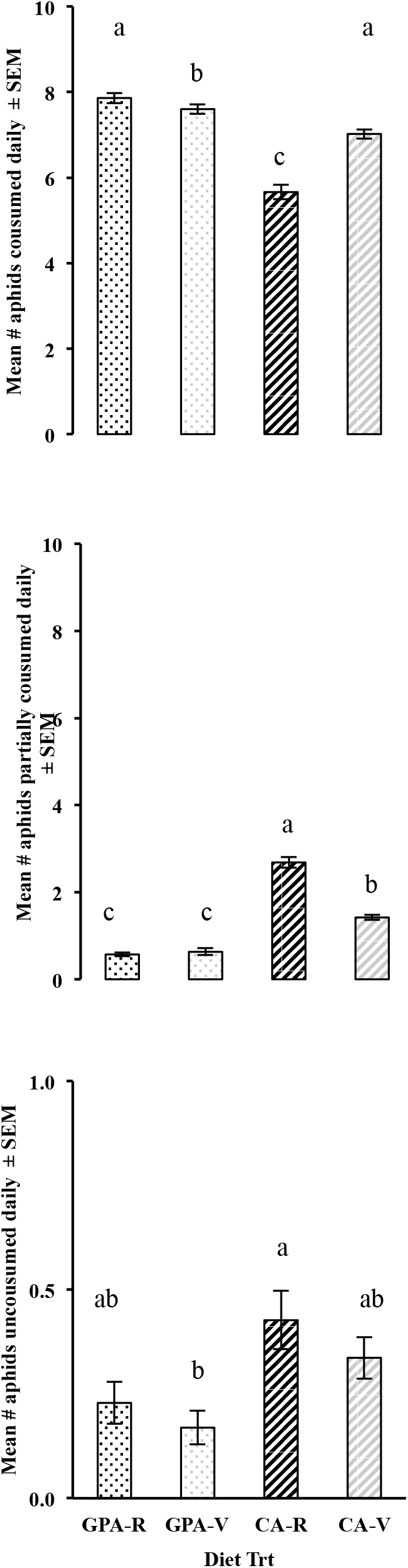
Mean aphid daily consumption ± SEM for *H. convergens* larvae restricted to different canola aphid diets: *Myzus persicae* (GPA) and *Brevicoryne brassicae* (CA) previously confined to reproductive (GPA-R and CA-R) or vegetative (GPA-V and CAV) structures of canola plants. Daily consumption was categorized as mean number of aphids (10 initial aphids) that were A) fully consumed, B) partially consumed, C) and unconsumed. Bars with the same letters are not significantly different (*P*< 0.05) using Fisher’s Protected LSD.

Differences in *H. convergens* consumption across diets were also compared among larval instars. Diet had a significant effect on consumption for all instars (first: *F*= 12.72; df= 4, 85; *P* <0.0001; second: *F*= 7.99; df= 4, 85; *P* <0.0001; third: *F*= 4.94; df= 4, 85; *P* =0.0013; fourth: *F*= 3.05; df= 4, 85; *P* =0.0213). First instars consumed the most SA, followed by GPA, and then CA regardless of previous feeding location by prey (Fig. 5). Second instars consumed significantly more SA than any other diet, but there were no differences in consumption among canola aphid species or feeding locations (Fig. 5). For the third instar, prey consumption was highest on the SA diet, followed by the GPA diets, and the CA diet regardless of previous feeding location, although CA-V had no statistical differences with either GPA diets and CA-R respectively (Fig. 5). Plant feeding location of aphids did not appear to affect third instar prey consumption. In contrast, prey consumption by fourth instars did not differ significantly among diets except for CA-R where larvae consumed 12% fewer prey than any of the other aphid diets (Fig. 5).

**Figure 5.**
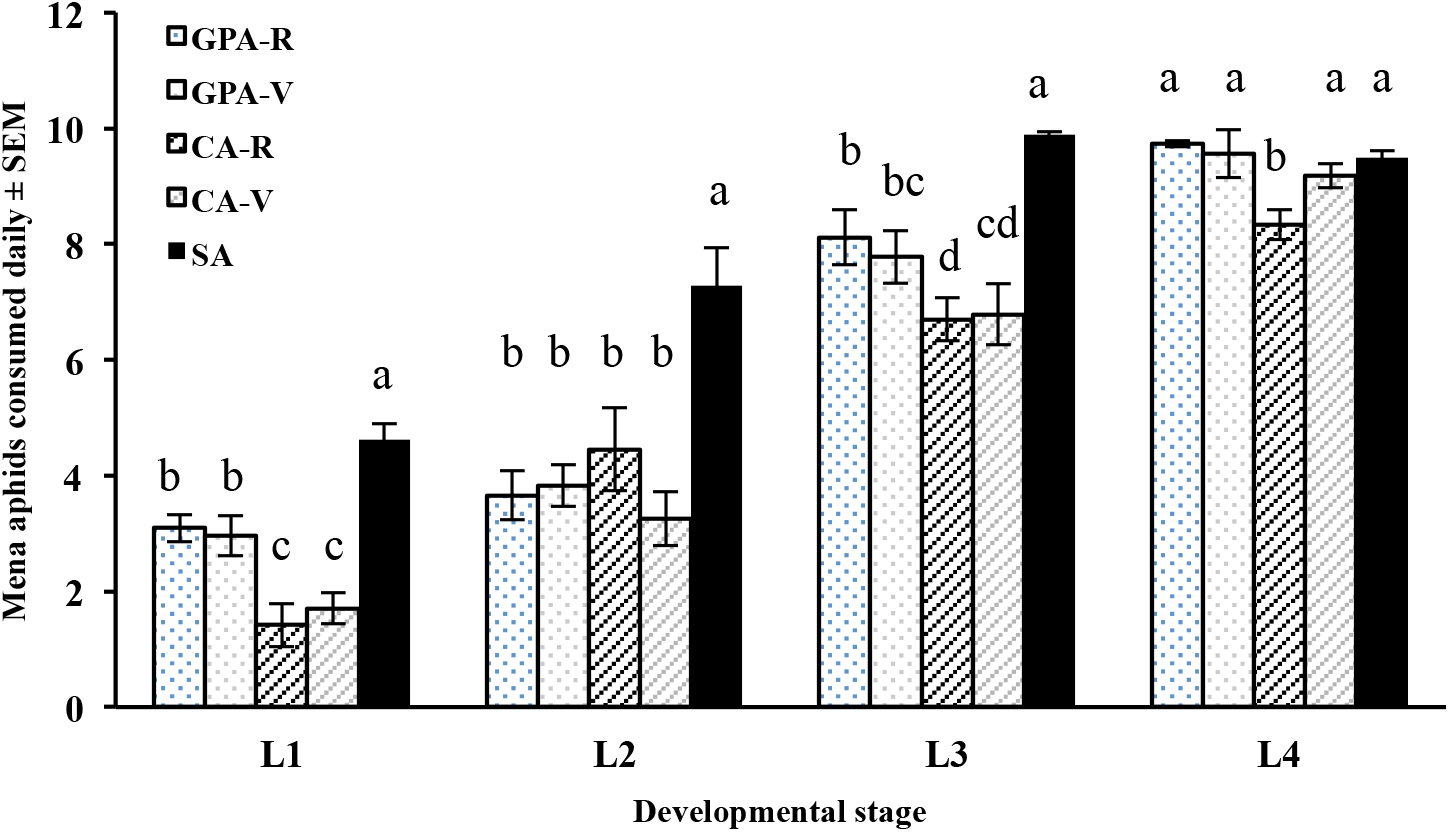
Mean number of aphids consumed ± SEM within each *H. convergens* larval stage for aphid diets tested: *Myzus persicae* (GPA) and *Brevicoryne brassicae* (CA) previously confined to reproductive (GPA-R and CA-R) or vegetative (GPA-V and CA-V) structures of canola plants, and an aphid control (*Aphis glycines*, SA). Bars with the same letters are not significantly different (*P*< 0.05) using Fisher’s Protected LSD.

### Pupal weight

There were significant differences in *H. convergens* pupal weights among diets treatments (*F*= 48.67, df= 5, 94; *P* <0.0001). The highest pupal weights were observed when lady beetle larvae consumed control diets, E-EGGS, followed by SA. Lowest weights occurred on canola aphid diets, regardless of treatment (GPA-R, GPA-V, CA-R, CA-V), in these where pupae treatment, where pupae weighed 40% less than those that fed on SA control aphid diets (Fig.6). When comparisons for pupal weight were made for canola aphid diet exclusively (CA-R, CA-V, GPA-R, GPA-V), only the main effect of aphid species (spp.) had a significant consequence on pupal weight; with pupa weighing 9.82 (± 0.29) g and 8.98 (± 0.25) g on CA and GPA, respectively (Table 1F).

**Figure 6.**
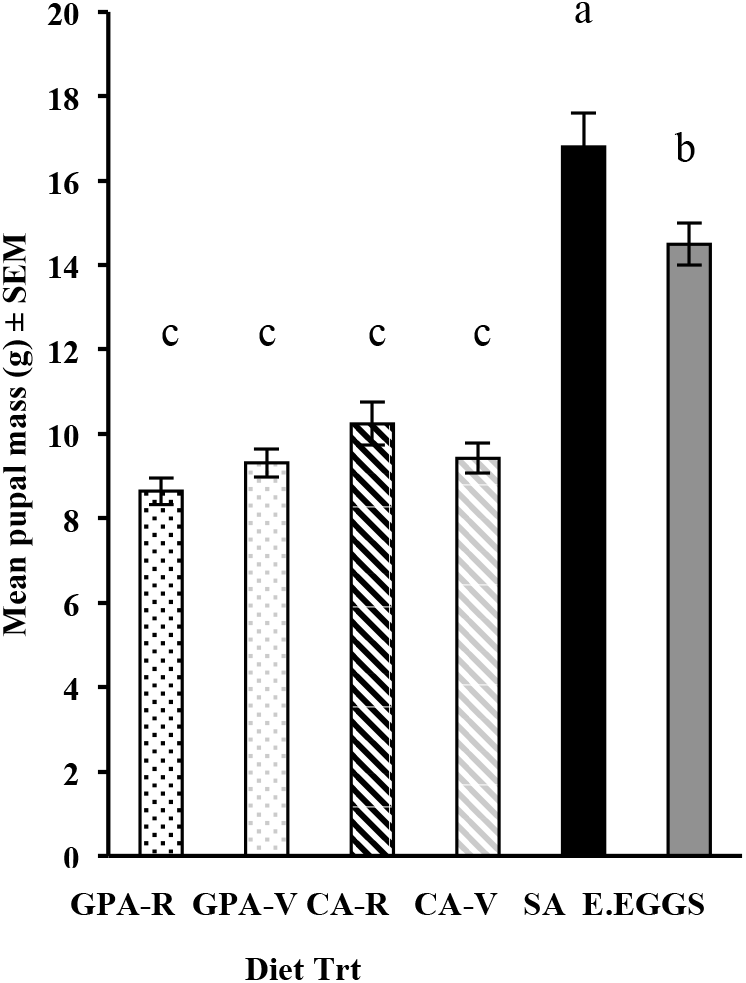
Mean pupal weights ± SEM for *H. convergens* larvae restricted to six different diets. Diet treatments included: *Myzus persicae* (GPA) and *Brevicoryne brassicae* (CA) previously confined to reproductive (GPA-R and CA-R) or vegetative (GPA-V and CA-V) structures of canola plants, an aphid control (*Aphis glycines*, SA), and a non-aphid diet (*Ephestia kuehniella* eggs). Bars with the same letters are not significantly different (*P*< 0.05) using Fisher’s Protected LSD.

## Discussion

Larval diet influenced life history responses of *H. convergens*, but the effects varied depending on the trait measured, which prey was offered (canola vs. no-canola aphid), and where on the plant they fed (reproductive vs. vegetative). Among aphid diets, *H. convergens* larvae had the longest development times on SA and CA regardless of feeding location, and GPA that had fed only on reproductive parts of canola plants. Development was shortest when predators were fed GPA from vegetative plant parts, and lastly the non-aphid diet (Fig 1A, C and Fig 2A-B). Explanations for differences in *H. convergens* development could be related to allelochemicals affecting digestibility or to differences in nutritional quality of the aphid diets (Jessie et al. 2015). Among canola aphid diets, differences in *H. convergens* development may be linked to differences in concentrations of glucosinolates ingested by larvae for several reasons. First, canola aphids have evolved different mechanism to process glucosinolates and other allelochemicals produced by canola plants. CA sequesters (retains) toxic compounds in their bodies, whereas GPA excretes these deleterious chemicals (Weber 1985). Second, higher concentrations of glucosinolates are found in reproductive tissues of canola plants compared to vegetative tissues (Cibils-Stewart 2013). This is consistent with our observation that total development of *H. convergens* was longest on GPA previously reared on reproductive parts of canola. Although we did not measure glucosinolate concentrations in aphids from different canola structures, we surmise that, even with its ability to excrete glucosinolates, concentrations in the GPA diet from reproductive tissues of canola may have been higher than those of GPA restricted to vegetative tissues; thus, this appears to influence *H. convergens* development (Cibils-Stewart 2013). Furthermore, no difference in overall larval development was observed between the two CA diets, suggesting that maximum sequestration of allelochemicals may have occurred, thus masking any effect due to plant feeding location. We would expect CA to have the highest glucosinolate concentrations, which could have had a combined effect on larval and pupal development; future studies that measure levels of glucosinolates in pupae will help quantify this effect. Our findings suggest that even with different mechanisms for processing glucosinolates, concentrations of GLS, or other allelochemicals, were higher in aphids taken from reproductive tissues of canola plants. Other studies focusing on various brassica herbivores have documented the negative effects of glucosinolates on natural enemy development as well as other fitness traits, further supporting our findings (Blackman 1967, Gabrys et al 1997, Cole. 1997a, Van Dam et al 2008a, Van Dam et al 2008b, Guigo et al. 2010, Kabouw et al 2010, Kramer et al. 2011).

In contrast to canola aphids, longer development of *H. convergens* larvae on soybean aphids may have been associated with low nutritional quality of this diet. For example, the consumption rate of soybean aphids was highest of all aphid diets, and was associated with the longest larval development times (Fig. 1A, C). This finding may indicate compensation for reduced *per capita* nutrient intake. Relationships between prey size, nutritional quality, and consumption can best be determined if experiments provide predetermined amounts of aphids on a biomass basis to control for intake rates and differences in aphid size across species (Jessie 2013). For example, other studies report differences in consumption rates that are related to prey size for other lady beetle species (Elliott et al. 1994, Tsaganou et al. 2004). Nutritional quality, such as proportion of lipids and/or protein of aphids from different plant structures should also be measured since it can directly influence predator performance (Giles et al. 2002).

Consumption of canola aphids was affected by aphid species and feeding location on host plants. *H. convergens* larvae consumed fewer CA than GPA. Of those that fed on *B. brassicae*, the fewest aphids were consumed, and more partially consumed and unconsumed aphids were left behind when aphids were obtained from reproductive canola structures (Fig 3A-C and Fig. 4A-C). For *H. convergens* feeding on canola aphids, we assumed that differences in development were related solely to levels of allelochemicals ingested by larvae (Cibils-Stewart et al. 2013). However, the amount of prey consumed can be affected by prey size, which is something we did not measure. Prey consumption may also interact with diet quality in at least two ways. First, if an unlimited number of aphids is provided, as was done in our study, predators may compensate for poor quality by increasing consumption. For example, Phoofolo et al. (2007) showed that the effect of nutritional quality of cereal aphids for *H. convergens* could only be measured when prey were provided at suboptimal levels. Second, an inferior diet may affect feeding behavior by reducing prey consumption. In our study, *H. convergens* consumed fewer canola aphids in situations where glucosinolates were likely at higher levels; we do not know if the apparent negative effect of these allelochemicals was direct, indirect, or both. It is also possible that prey consumption was influenced by prey size (not measured in this study) or nutritional quality unrelated to allelochemicals (e.g., sugars, fatty acids, etc.). When compared to canola aphids, consumption of soybean aphids was much greater, and partially consumed and unconsumed aphids lower, suggesting either that toxicity of SA was lower, that they were nutritionally inferior, or a combination (Fig 3A-C). When considering the effect of diet consumption in each instar, it seems that canola aphid diets had a greater effect on smaller larvae, where *H. convergens* consumed fewer glucosinolate-sequestering aphids (CA) than glucosinolate-excreting (GPA) aphids compared to SA. Our results are consistent with Fisker and Tolf (2004) that indicated ontogenetically increased tolerance when more mature spiders were fed toxic pray compared to younger instar exposure. This suggests a relationship between predator size and prey toxicity exists in another predator/prey system.

Understanding canola-aphid-predator interactions has important implications for pest management. Canola producers currently manage aphid outbreaks by applying pesticides during the canola blooming period. Current monitoring programs do not record aphid distributions within the canola canopy or potential effects of glucosinolate sequestration on natural enemy communities. Understanding the interactions between secondary compounds, feeding location, and its relation to allelochemicals sequestration by aphids and potential impacts on the natural enemy community might facilitate best management practices that reduce pesticide usage and enhance natural control in winter canola fields.

Understanding aphid demography in canola, and interactions with natural enemies might improve monitoring and management programs, and help to understand effects of allelochemicals at other trophic levels, such as effects of prey feeding location on natural enemy communities. Our results indicate that feeding location influences demography of aphids directly, different plant structures are being either sources or sinks to CA. Source-sink relationships within the plant are directly affecting demography of aphids within the canopy and therefore creating additional source-sinks within the canopy for third tropic levels. Varying degrees of allelochemicals acquisition by the cabbage aphids in different plant structures may be directly influenced by feeding location, which is a new contribution to agro-toxicology and ecotoxicology. By evaluating source-sink dynamics at the plant level, we can further understand trophic interactions occurring between species (both pest and beneficial) at the landscape-level. More specifically, such interactions may influence community dynamics and structure for newly introduced crops into established agroecosystems (Guigo and Corff 2010)

Overall, our data indicate that there is considerable variation in life history responses of *H. convergens* on different diets. Larvae are capable of surviving exclusively on a diet of canola aphids, although feeding location within a plant directly affects their development rate. These trade-offs between prey feeding locations are important to understand, especially in the context of adding new crops to an agricultural landscape and understanding the implications of such actions on existing natural enemy communities. Our results indicate that interactions between predator fitness and aphid feeding location are likely to be encountered in the field and may result in negative repercussions for predator life history traits, as observed in our study. Future studies should examine effects of diet on *H. convergens* by evaluating daily consumption in terms of grams of prey, not numbers, duration of the feeding period, and weight changes for each larval instar (Soares et al. 2001). In addition, prey size should be recorded since studies with other lady beetle species have reported differences in consumption rates related to prey size (Elliott et al. 1994, Tsaganou et al. 2004). To assess prey quality, it would be useful to provide lady beetles with predetermined weights of aphids when measuring the amount of prey consumed. Finally, both nutritional (fatty acids, protein, total caloric content, etc.) and allelochemical data should be collected and compared from different plant structures (and from the aphid prey) with respect to predator performance (Giles et al. 2002). Field studies to determine recruitment, oviposition, and feeding behavior of *H. convergens* adults on different canola aphids previously reared on either reproductive or vegetative parts of the canola plant are needed as they would help to understand how plant and aphid chemistry shape predation and, thus, biological control of canola aphid populations. In addition, since predators often encounter different prey species simultaneously, choice studies involving mixed species diets would further our understanding of choices *H. convergens* makes in the canola system under more realistic field situations. Finally, behavioral studies that analyze coccinellid feeding behavior under these complex environmental conditions will better enable an understanding of predator movement among and between crops, ultimately resulting in more predictive economic threshold models for aphid management in existing or newly introduced crop systems (Obrycki et al. 2009).

## Acknowledgements

Our project was part of a collaboration between research scientists at Kansas State University, Oklahoma State University, University of Arkansas, and USDA ARS in Arizona. Canola seeds used to rear aphid colonies were provided by M. Stamm canola breeder with the department of Agronomy at Kansas State University. Thanks to the McCornack Lab field crew for assistance with field and greenhouse project and to the Department of Entomology at Kansas State University. Financial support for this research was partially provided by the USDA-NIFA AFRI (# 33488) Sustainable Biofuels Initiative and the Kansas Agricultural Experiment Station (KAES); this article is contribution 19–047-J from the KAES.

## References

Baggen, L.R., G.M. Gurr, and A. Meats. 1999. Flowers in tri-trophic systems: mechanisms allowing selective exploitation by insect natural enemies for conservation biological control. Entomologia Experimentalis et Applicata 91: 155–161.

Berberet, R.C., K.L. Giles, A.A. Zarrabi, and M.E. Payton. 2009. Development, reproduction, and within-plant infestation patterns of *Aphis craccivora* (Homoptera: Aphididae) on alfalfa. Environmental Entomology 38: 1765–1771.

Berlandier, F., D. Severtson and P. Mangano. 2010. Aphid management in canola crops. Department of Agriculture and Food, Government of Western Australia. Fact sheet ISSN 0726–934X.

Berner, D., and W.U. Blanckenhorn. 2007. An ontogenetic perspective on the relationship between age and size at maturity. Functional Ecology 21:505–512.

Blackman, R.L. 1967. The effects of aphid foods on *Adalia bipunctata* Linnaeus and *Cocciniella septempunctata* Linneus. Annuals Applied Biology 59: 207–219.

Blackshaw, R.E., F.J. Larney, C.W. Linswall, P.R. Watson, and D.A. Derkson. 2001. Tillage intensity and crop rotation affect weed community dynamics in winter wheat cropping systems. Canadian Journal of Plant Science 81: 805–813.

Brewer, M.J., and N. C. Elliott. 2004. Biological control of cereal aphids in North America and mediating effects of host plant and habitat manipulation. Annual Review of Entomology 49: 219–242.

Brown, P.D., J.G. Tokuhisa, M. Reichelt, and J. Gershenzon. 2003. Variation of glucosinolate accumulation among different organs and developmental stages of *Arabidopsis thaliana*. Phytochemistry 62: 471–481.

Cabral, S., A.O. Soares, R. Moura, and P. Garcia. 2006. Suitability of *Aphid fabae, Myzus persicae* (Homoptera: Aphididae) and *Aleyrodes proletella* (Homoptera: Aleyrodidae) as prey for *Coccinella undecimpuntata* (Coleoptera: Coccinellidae). Biological Control 39: 343–440.

Chambers, R.J. 1986. Control of cereal aphids in winter wheat by natural enemies: aphid-specific predators, parasitoids and pathogenic fungi. Annals of Applied Biology 108: 219–231.

Cibils-Stewart, X. 2013. Influence of Plant Architecture on Tritrophic Interactions between Winter Canola (Brassicae napus), Aphids (Hemiptera: Aphididae) and *Hippodamia convergens* (Coleoptera: Coccinellidae). MSc Thesis, Kansas State University, Manhattan, KS, USA. Available at: hdl.handle.net/2097/16875

Ciblis-Stewart X., B.P. McCornack and B. Sandercock. 2015. Feeding location affects demographic performance of cabbage aphids on winter canola. Entomologia Experimentalis et Applicata. 156(2):149–159.DOI: 10.1111/eea.12325.

Cole, R.A. 1997a. Comparison of feeding behavior of two brassica pests *Brevicoryne brassicae* and *Myzus persicae* on wild and cultivated brassica species. Entomologia Experimentalis et Applicata 85: 135–143.

Cole, R.A. 1997b. The relative importance of glucosinolates and amino acids to the development of two aphid pests *Brevicoryne brassicae* and *Myzus persicae* on wild and cultivated brassica species. Entomologia Experimentalis et Applicata 85: 121–133.

Dixon, A.F.G. 2000. Insect Predator–Prey Dynamics: Ladybird Beetles and Biological Control. Cambridge University Press, Cambridge, UK. 258 pp.

Dogan, E.B., R.E. Berry, G.L. Reed, and P.A. Rossignol. 1996. Biological parameters of convergent lady beetles (Coleptera: Coccinellidae) feeding on aphids (Homoptera: Aphididae) on transgenic potato. Biological and Microbial Control 89: 1105–1108.

Dwernychuk, L.W., and D.A Boag. 1972. Ducks nesting in association with gulls-an ecological trap?. Canadian Journal of Zoology 50: 559–563.

Elliott, N.C., B.W. French, G. L. Michels, and D.K. Reed. 1994. Influence of four aphid species on development, survival and adult size of *Cyclonedamunda* (Gremar) (Coleoptera: Coccinellidae). Southwestern Entomologist 19:57–61.

Evans, E.W. 2003. Searching and reproductive behavior of female aphidophagous ladybirds (Coleoptera: Coccinellidae): a review. European Journal of Entomology 100: 1–10.

Fehr, W.R., C.E. Caviness, D.T. Burmood, and J.S. Pennington. 1971. Stage of development descriptions for soybeans, *Glycine max* (L.) Merr. Crop Science 11:929–931.

Fernandes, F.S., R.S. Ramalho, J.L. Nascimento Junior, J.B. Malaquias, A.B.B. Nascimento, C.A.D. Silva, and J.C. Zanuncio. 2011. Within-plant distribution of cotton aphids, *Aphis gossypii* Glover (Hemiptera: Aphididae), in Bt and non-Bt cotton fields. Bulletin of Entomological Research 102: 79–87.

Fisker, E., and Tolf S. 2004. Effects of chronic exposure to prey in generalist predator. Physiological Entomology 29 (2):129–138.

Francis, F., E. Haubruge, and C. Gaspar. 2000. Influence of host plant specialist/generalist aphids and the development of *Adelia bipunctata* (Coleoptera: Coccinellidae). European Journal of Entomology 97: 481–485.

Francis, F., G. Lognay, J-P. Wathelet, and E. Haubruge. 2001. Effects of allelochemicals from first (brassicae) and second (*Myzus persicae* and *Brevicoryne brassicae)* trophic levels on *Adelia bipunctata*. Journal of Chemical Ecology 27: 243–56.

Franke, T.C., K.D. Kelsey, and T.A. Royer. 2009. Pest management needs assessment for Oklahoma canola producers. Oklahoma Cooperative Extension Service EPP-7085.

Gabrys, B., W.F. Tjallingii, and T.A. Van Beek. 1997. Analysis of probing by *Brevicoryne brassicae* on host plant parts with different glucosinolate contents. Journal of Chemical Ecology. 23: 1661–1673.

Giles and Walker. 2009. Chapter 16: Dissemination and Impact of IPM programs in US agriculture. Integrated Pest Management: Dissemination and Impact Springer editions.

Giles, K., G. Hein, and F. Peairs. 2008. Chapter 19: Areawide pest management of cereal aphids in dryland wheat systems of the great plains, USA. Areawide Pest Management: Theory and Implementation, Opender, Koul, Gerrit W. Cuperus, and Norman Elliott (Eds.). CAB International.

Giles, K.L., J.W. Dillwith, R.C. Berberet, and N.C. Elliott. 2005. Survival, development, and growth of *Coccinella septempunctata* fed *Schizaphis graminum* from resistant and susceptible cultivars. Southwestern Entomologist 30: 113–120.

Giles, K.L., R.D. Madden, R. Stockland, M.E. Payton, and J.W. Dillwith. 2002. Host plants affects predator fitness via the nutritional value of herbivore prey: Investigation of a plant-aphid-lady beetles system. BioControl 47: 1–21.

Guigo, P.L., Y. Qu, and J.L. Corff. 2010. Plant-mediated effects on a toxin-sequestering aphid and its endoparasitoid. Basic and Applied Ecology 12: 72–79.

Hajek, A. 2004. Natural enemies: An introduction to biological control. Cambridge University Press.

Hodek, I. 1973. Biology of Coccinellidae. Dr. W. Junk. N.V., Publishers, the Hague.

Hodek, I., and A. Honěk. 1996. Ecology of Coccinellidae. Kluwer Academic, Dordrecht.

Hopkins, R. J., N. M. Van Dam., and J. J.A. Van Loon. 2009. Role of glucosinolates in insect-plant relationships and multitrophic interactions. Annual Review of Entomology 54: 57–83.

Hughes, R.D. 1963. Population dynamics of the Brevicoryne brassicae, *Brevicoryne brassicae* (L.). Journal of Animal Ecology 32:393–424.

Idris, A.B. and M.N.M. Roff. 2002. Vertical and temporal distribution of *Aphis gossypii* Glover and coccinellid populations on different chilli (*Capsicum annum*) varieties. Journal of Asia-Pacific Entomology 5: 185–191.

Jeger, M.J., and S.L.H. Viljinen-Rollinson. 2001. The use of the area under the disease-progress curve (AUDPC) to assess quantitative disease resistance in crop cultivars. Theoretical and Applied Genetics 102(1):32–40 · January 2001.

Jessie, C.N, 2017. Diversity, relative abundance, movement, and fitness of insect predators in winter canola agricultural landscapes in the United States southern plains. Ph.D Dissertation, Oklahoma State University.

Jessie, W. P., K. L. Giles, E. J. Rebekj, M. E. Payton, C. N. Jessie and B. P. Mccornack. 2015. Preference and Performance of *Hippodamia convergens* (Coleoptera: Coccinellidae) and *Chrysoperla carnea* (Neuroptera: Chrysopidae) on *Brevicoryne brassicae, Lipaphis erysimi*, and *Myzus persicae* (Hemiptera: Aphididae) From Winter-Adapted Canola. Environmental Entomology. 44: 880–889.

Kabouw, P., A. Biere, W. H. Van Der Putten, and N. M. Van Dam. 2010. Intraspecific differences in root and shoot glucosinolate profiles among white cabbage (*Brassica oleracea* var. capitata) cultivars. Journal of Agricultural and Food Chemistry 58: 411–417.

Kajita, Y., and E.W. Evans. 2010. Relationship of body size, fecundity, and invasion success among predatory lady beetles (Coleoptera: Coccinellidae) inhabiting alfalfa fields. Annuals of the Entomology Society of America 103:750–756.

Kalushkov, P., and I. Hodek. 2001. New essential aphid prey for *Anatis ocellata* and *Calvia quatuordecimguttata* (Coleoptera: Coccinellidae). BiocontrolScience and Technology 11: 35–39.

Kazana, E., T.W. Pope., L. Tibbles., M. Bridges., J. A. Pickett., A. M. Bones., G. Powell., and J.T. Rossiter. 2007. The Brevicoryne brassicae: a walking mustard oil bomb. Proceedings of the Royal Society Biological Sciences 274, 2271–2277.

Knodel J.J. 2011. Scout for aphids in canola http://www.ag.ndsu.edu/cpr/entomology/Nscout-for-aphids-in-canola-7–28–11

Kos, M., C. Broekgaarden, P. Kabouw, K.O. Lenferink, E.H. Poelman, L.E.M. Vet, M. Dicke, and J.J.A Van Loon. 2011a. Relative importance of plant-mediated bottom-up and top-down forces on herbivore abundance on *Brassica oleracea*. Functional Ecology 25: 1113–1124.

Kos, M., P. Kabouw., R. A. Noordam., K. Hendriks., L. E. M. Vet., J. J. A. Van Loon., and M. Dicke. 2011b. Prey-mediated effects of glucosinolates on aphid predators. Ecological Entomology 36, 377–388.

Kova’r, I. 1996. Morphology and Anatomy. Pp:1–18. In: I. Hodek and A. Honek, eds. Ecology of Cocinellidae. Kluwer Academic Publishers.

Kramer R.C., D.J. Kliebenstein, A. Chiem, E. Morrill, N. J. Mills, and C. Kremen. 2011. Chemically mediated tritrophic interactions: opposing effects of glucosinolates on a specialist herbivore and its predators. Journal of Applied Ecology 48:880–887.

Lambton, P.W., and M. Hassall. 2005. How should toxic secondary metabolites be distributed between the leaves of a fast-growing plant to minimize the effects of herbivory? Functional Ecology 19: 299–305.

Landis, D.A., S.D. Wratten, and G. M. Gurr. 2008. Habitat management to conserve natural enemies of arthropod pests in agriculture. Annual Review of Entomology 45: 175–201.

Lee, J-H., and T-J. Kang. 2004. Functional response of *Harmonia axyridis* (Pallas) (Coleoptera: Coccinellidae) to *Aphis gossipii* Glover (Homoptera: Aphididae) in the laboratory. Biological Control 31: 306–310.

Lu, Y., K. Wu, Y. Jiang, Y. Guo and N. Desneux. 2012. Widespread adoption of bt cotton and insecticide decrease promotes biocontrol services. Nature 487.7407: 362–5.

Lundgren, J.G., S.E. Moser, R.L. Hellmich and M.P. Seagraves. 2011. The effects of diet on herbivory by a predaceous lady beetle. Biocontrol Science and Technology, 21: 71–74. doi:10.1080/09583157.2010.524917, First posted on: 24 September 2010.

Merritt, S.Z. 1996. Within-plant variation in concentrations of amino acids, sugar, and sinigrin on phloem sap of black mustard, *brassica nigra* (L.) Koch (Crusiferae). Journal of Chemical Ecology 22: 1133–1145.

Michaud, J.P., and J.A. Qureshi. 2006, Reproductive diapause in *Hippodamia convergens* (Coleoptera: Coccinellidae) and its life history consequences. Biological Control 39: 193–200.

Michels G.L., N.C. Elliott, R.A. Romero, D.A. Owings, and J.B. Bible. 2001. Impact 0f indigenous lady beetles on Russian wheat aphids and greenbugs (Homoptera: Aphididae) infesting winter wheat in the Texas panhandle. Southwestern Entomologist 26:77–113.

Michels, G.J. and R. W. Behle. 1991. A comparison *Coccinella septempunctata* and *Hippodamia convergens* larval development on greenbugs at constant temperatures. Southwestern Entomologist 16:73–80.

Murphy, L. A., and R. Scarth. 1994. Vernalization response in spring oilseed rape (*Brassica napus* L.) cultivars. Canadian Journal of Plant Science 74: 275–277.

Nechols, J. R. and T. L. Harvey. 1998. Evaluation of a mechanical exclusion method to assess the impact of Russian wheat aphid (Homoptera: Aphididae) natural enemies. Pp. 270–279. *In* S. S. Quisenberry and F. Peairs(eds.), Response Model for an Introduced Pest--The Russian Wheat Aphid (Homoptera: Aphididae). Thomas Say Publications in Entomology, Entomological Society of America, Lanham MD.

Nielson, M.W. and W.E. Currie.1959. Biology of the convergent lady beetles when fed a spotted alfalfa aphid diet. Journal Economic Entomology 53 (2): 257–259.

Obrycki, J.J., and T.J. Kring. 1998. Predaceous Coccinellidae in biological control. Annu. Rev. Entomol. 43: 295–321.

Obrycki, J.J., J.D. Harwood, T.J. Kring, and R.J. O’Neil. 2009. Aphidophagy by Coccinellidae: Applications of biological control in agroecosystems. Biological Control 51:244–254.

Patt, J. M., G.C. Hamilton, and J.H. Lashomb. 1997. Foraging success of parasitoid wasps on flowers: interplay of insect morphology, floral architecture and searching behavior. EntomologiaExperimentalisetApplicata 83: 21–30.

Peeper, T., H. Sanders, M. Boyles, and R. Gribble. 2009. OKANOLA. OAES and OCES Oklahoma State University. http://www.canola.okstate.edu/index.htm

Pekar, S. 2005. Horizontal and vertical distribution of spiders (Araneae) in sunflowers. Journal of Acarology 33: 197–204.

Perterson, G.A., and D.G. Westfall. 1994. Intensified cropping systems: the key to environmental and economic sustainability in the Great Plains. In Proceedings of the intensive wheat management conference, Norcoss, Gerogia, Potash and Phosphate institute, pp. 73–84.

Phoofolo, M.W., K.L. Giles, and N.C. Elliott. 2007. Quantitate evaluation of suitability of the greenbug, *Schizaphis gramunum*, and the bird cherry-oat aphid, *Rhopalosiphum padi*, as prey for *Hippodamia convergens* (Coloeoptera: Coccinellidae). Biological Control 41:25–32.

Phoofolo, M.W., K.L. Giles, and N.C. Elliott. 2008. Larval life history responses to food deprivation in three species of predatory lady beetles (Coleoptera: Coccinellidae). Environmental Entomology 37: 315–322.

Phoofolo, M.W., N.C. Elliott, and K.L. Giles. 2009. Analysis of growth and development in the final instar of three species of predatory Coccinellidae under varying prey availability. Entomologia Experimentalis et Applicata 131: 264–277.

Pratt, C., T.W. Pope., G. Powell., and J.T. Rossiter. 2008. Accumulation of glucosinolates by the Brevicoryne brassicae *Brevicoryne brassicae* as a defense against two coccinellid species. Journal of Chemical Ecology 34: 323–329.

Royer, T.A. and K. Giles. 2008 (rev. 2010).Management of insect and mite pests in canola. Oklahoma Cooperative Extension Service Fact Sheet CR-7667.

Royer T.A. and K.L. Giles. 2017. The OKANOLA project: challenges in managing insect pests of canola in the southern plains. In: Reddy, G.V.P. Integrated Management of Insect Pests on Canola and Other Brassica Oilseed Crops. CAB International, Boston, MA. 147–156.

Seagraves, M.P. 2009. Lady beetle oviposition behavior in response to the trophic environment. Biological Control 51: 313–322.

Smallengange, R.C., J. J. A. Van Loon, S. E. Blatt, J. A. Harvey, N. Agerbirk, and M. Dicke. 2007. Flowers vs. leaf feeding by *Prieris brassicae:* Glucosinolate-rich flower tissues are preferred and sustain higher growth rates. Journal of Chemical Ecology 33: 1831–44.

Soares, A.O., D. Coderre, and H. Schanderl. 2001. Fitness of two phenotypes of *Harmonia axyridis* (Coleoptera: Cocinellidae). European Journal of Entomology 98:287–293.

Soper, A.M., R.J. Whitworth, and B.P. McCornack. 2013. Sorghum seed maturity affects weight and feeding duration of immature *Helicoverpa zea* and *Spodoptera fungiperda* (Lepidoptera: Noctuidae) in the laboratory. Journal of Insect Science 13: 67.

Tickell, J. 2000. From the fryer to the fuel tank: the complete guide to using vegetable oil as an alternative fuel. 3rd edition. Tickell Energy Consulting, Tallahassee, FL. 162 p.

Tsaganou, F.C., C.J. Hodgson, C.G. Athanassiou, N.G. Kavallieratos, and Z. Tomanovic’. 2004. Effects of *Aphis gossipii* Glover, *Brevicoryne brassicae* (L.), and *Megoura viciae* Buckton (Hemiptera:Aphidoidea) on the development of the predator *Harmonia axyridis* (Pallas) (Coleoptera: Coccinellidae). Biological Control 31: 138–144.

U.S. Department of Agriculture.2012 Crop Production Report. https://www.usda.gov/media/agency-reports (Accesses: July 2018).

US Canola Association website. http://www.uscanola.com/ (Accesses: July 2018).

Van Dam, N.M., and M. W. A. T. Oomen. 2008a. Root and shoot jasmoinc acid applications differentially affect leaf chemistry and herbivore growth. Plant Signaling and Behavior 3: 91–98.

Van Dam, N.M., T. O. G. Tytgat, and J. Kirkegaard. 2008b. Root and shoot glucosinolates: a comparison of their diversity, function, and interactions in natural and managed ecosystems. Phytochemistry Reviews 8: 171–186.

Vargas, G., J.P. Michaud, and J.R. Nechols. 2012. Larval supply constrains female reproductive schedules in *Hippodamia convergens* (Coleoptera: Coccinellidae). Annuals of the Entomology Society of America 105: 832–839.

Weber, G. 1985. Genetic variability in host plant adaptation of the Myzus persicae, *Myzus persicae*. Entomologia Experimentalis et Applicata 38: 49–56.

Wolfenbarger, L. L., S. E. Naranjo, J. G. Lundgren, R. J. Bitzer, and L. S. Watrud. 2008. Bt crop effects on functional guilds of non-target arthropods: a meta-analysis. PLoS ONE 3: e2118. doi:10.1371/journal.pone.0002118.

